# Characterization and simulation of metagenomic nanopore sequencing data with Meta-NanoSim

**DOI:** 10.1101/2021.11.19.469328

**Authors:** Chen Yang, Theodora Lo, Ka Ming Nip, Saber Hafezqorani, Rene L Warren, Inanc Birol

## Abstract

Nanopore sequencing is crucial to metagenomic studies as its kilobase-long reads can contribute to resolving genomic structural differences among microbes. However, platform-specific challenges, including high base-call error rate, non-uniform read lengths, and the presence of chimeric artifacts, necessitate specifically designed analytical tools. Here, we present Meta-NanoSim, a fast and versatile utility that characterizes and simulates the unique properties of nanopore metagenomic reads. Further, Meta-NanoSim improves upon state-of-the-art methods on microbial abundance estimation through a base-level quantification algorithm. We demonstrate that Meta-NanoSim simulated data can facilitate the development of metagenomic algorithms and guide experimental design through a metagenomic assembly benchmarking task.

## BACKGROUND

Empowered by the rapid development of next-generation sequencing technologies, metagenomic analysis has enabled comprehensive investigation of the genetic composition and abundance of microbial communities. In investigating microbiomes, metagenomic sequencing bypasses the need to culture each individual species by extracting DNA directly from their natural habitat, making it feasible to study the microbes that cannot be isolated or cultured in laboratory (1,2). Within the past few decades, the improved throughput and reduced cost of next-generation sequencing platforms have enabled a wide range of metagenomic studies of environmental, pharmaceutical, and medical relevance (3–5).

Until recently, Illumina short-read sequencing (Illumina Inc., San Diego, CA) has been the technology of choice for metagenomic sequencing projects due to its high throughput, low cost, and low error rate. However, the reads generated by Illumina instruments are often too short (<250 bp) to span inter- and intra-chromosomal homologous regions and suffer from intrinsic biases, thus complicating downstream assembly and taxonomic analysis (6). As a third-generation long-read sequencing technology from Oxford Nanopore Technologies Ltd. (ONT, Oxford, UK), nanopore sequencing is gaining traction in metagenomic research efforts, due largely to the long read lengths it generates, as well as the portability of their MinION sequencing platform (7). The N50 (the shortest read length to be included for covering 50% of the total sequenced length) in a typical run is over 5 kb (8) and the reported maximum read length exceeds 2 Mb. At the high end, whole bacterial or viral genomes may be captured by a few sequencing reads (9,10), making it possible to disambiguate between even closely-related strains. Since it was first announced, metagenomic sequencing by ONT has been playing an essential role in real-time pathogen identification and clinical diagnosis, including during the recent coronavirus disease 2019 pandemic (11–14).

Although a plethora of metagenomic analysis tools have been developed for short-read sequencing data, the challenges associated with ONT reads, such as high error rate, non-uniform error distributions, and chimeric read artifacts (8,15–17), are calling for analytical tools designed specifically for long reads. For example, quantification of microbial abundance levels, or metagenomic abundance estimation, is traditionally computed by counting the number of mapped reads followed by fine-tuning of ambiguous mappings (18,19). This approach has been proven to be cost-effective for Illumina short reads because of their uniform lengths. However, the accuracy of these tools would be understandably impacted when applied on ONT reads, especially for lowly represented genomes, because of the variable lengths and relatively high error rates compared to Illumina reads (5 - 15% depending on the flowcell chemistry and basecalling algorithm). In addition, ONT sequencing projects on genome, transcriptome, and metagenome, from prokaryotes to eukaryotes, were all reported to have certain problematic reads with gapped or chimeric alignments, likely generated due to library preparation or sequencing artifacts (17,20–25). Reference-based abundance estimation using merely primary alignments may further be affected by the presence of these chimeric reads, as well as reads that span the start position of a circular genome. To the best of our knowledge, even the state-of-the-art program, MetaMaps, does not account for chimeric reads, but simply uses an Expectation-Maximization (EM) algorithm to disambiguate multi-mapped reads (26). In this work, we show that there is still room for improving metagenomic abundance estimation, a proposition attainable by quantifying aligned bases instead of reads, while leveraging chimeric read information.

In the process of tool development and benchmarking, a metagenomic ONT read simulator and associated simulated datasets with known ground truth can save time and money. Ideally, such a read simulator should reflect the true characteristics of the ONT platform and allow effective evaluation of bioinformatics tools. In return, the evaluation results can guide the experimental design of metagenomics project, such as sequencing depths and number of replicates (27). Currently, the only simulator that specifically simulates ONT metagenomic datasets is CAMISIM (28). The workflow of CAMISIM is focused on the composition design of microbial community given a taxonomy profile. It uses NanoSim (version 1) (15) as the engine to simulate ONT reads for each genome separately once the composition of the community is determined. Following the same idea, one can also use other existing ONT genomic simulators naively to simulate each composite genome separately and then aggregate the reads according to the desired abundance. However, it is impractical to simulate a large microbial community with hundreds or more genomes with this approach, not to mention the existing simulators for ONT reads are not designed to model metagenomic specific features, such as chimeric reads and deviations in abundance levels. More importantly, the simulation of abundance levels should be consistent with the quantification method, thus merely mixing the reads from different genomes will yield a compromised abundance profile. Taken together, we believe the current version of NanoSim can be upgraded to capture and simulate read properties specific to metagenomics, especially the microbial abundance levels and chimeric reads – two key factors that may influence metagenome assembly, taxonomy binning, and abundance estimation. Further, in real world scenarios, viruses, bacteria, and fungi co-exist in complex microbial communities, hence it is a desired feature to simulate metagenomes comprising both circular and linear genomes.

Here, we introduce Meta-NanoSim (NanoSim version 3), an ONT metagenome simulator for complex microbial communities. Given a training dataset, Meta-NanoSim characterizes read length distributions, error profiles, and alignment ratio models. Optionally, it also detects chimeric reads and estimates microbial abundance levels. In our benchmarks, the performance of the metagenomic abundance estimation feature of Meta-NanoSim surpasses the current state-of-the-art methods. The chimeric read detection feature also improves upon the read length modelling in NanoSim v2, and thus simulating this artifact of the technology may challenge metagenomic analytical tools with a real-world scenario. Through benchmarking experiments comparing simulated reads with empirical datasets, we show that Meta-NanoSim preserves the key characteristics of ONT metagenomic reads. Further, we showcase the usability and utility of Meta-NanoSim with a metagenomic assembly task.

## RESULTS

Here, we first present the design of Meta-NanoSim and demonstrate the utility and performance of two key features, chimeric read detection and abundance estimation. To show the similarity between simulated reads and experimental reads, we generated a simulated dataset using models learned from experimental data and compared against a CAMISIM simulated dataset. We also show that Meta-NanoSim is capable of simulating large complex microbial community by simulating a dataset with 125 species with abundance levels estimated from a human saliva sample. Finally, we showcase a potential use case of Meta-NanoSim with a set of simulated data, benchmarking the metagenomic assembler MetaFlye (29) on how it scales with increasing sequencing depth. The input experimental data used in this section are two sets of publicly available mock community ONT sequencing reads, each dataset containing the same 10 microbial species (eight bacteria and two fungi) but with different abundance distributions. In this work, we denote the dataset with evenly distributed abundance levels as the *Even* dataset and the one with logarithmically distributed abundance levels as the *Log* dataset (detailed in Methods).

### Meta-NanoSim design

Meta-NanoSim is implemented in Python as two sub-modules in the NanoSim suite: `meta` in characterization and simulation stages. It learns the technical and metagenomic-specific features of ONT reads in the characterization stage, builds statistical models, and applies them in the simulation stage (Fig. 1). In the characterization stage, it takes ONT metagenomic reads and a reference metagenome as input to infer the ground truth through sequence alignments. Based on those alignments, it models the read length distributions (aligned and unaligned part) via kernel density estimation and basecall events via mixture statistical models. In addition to existing NanoSim features, we introduce two new analyses in its characterization pipeline; chimeric reads analysis for genome/metagenomes and abundance estimation for metagenomic datasets (Methods).

**Fig. 1.**
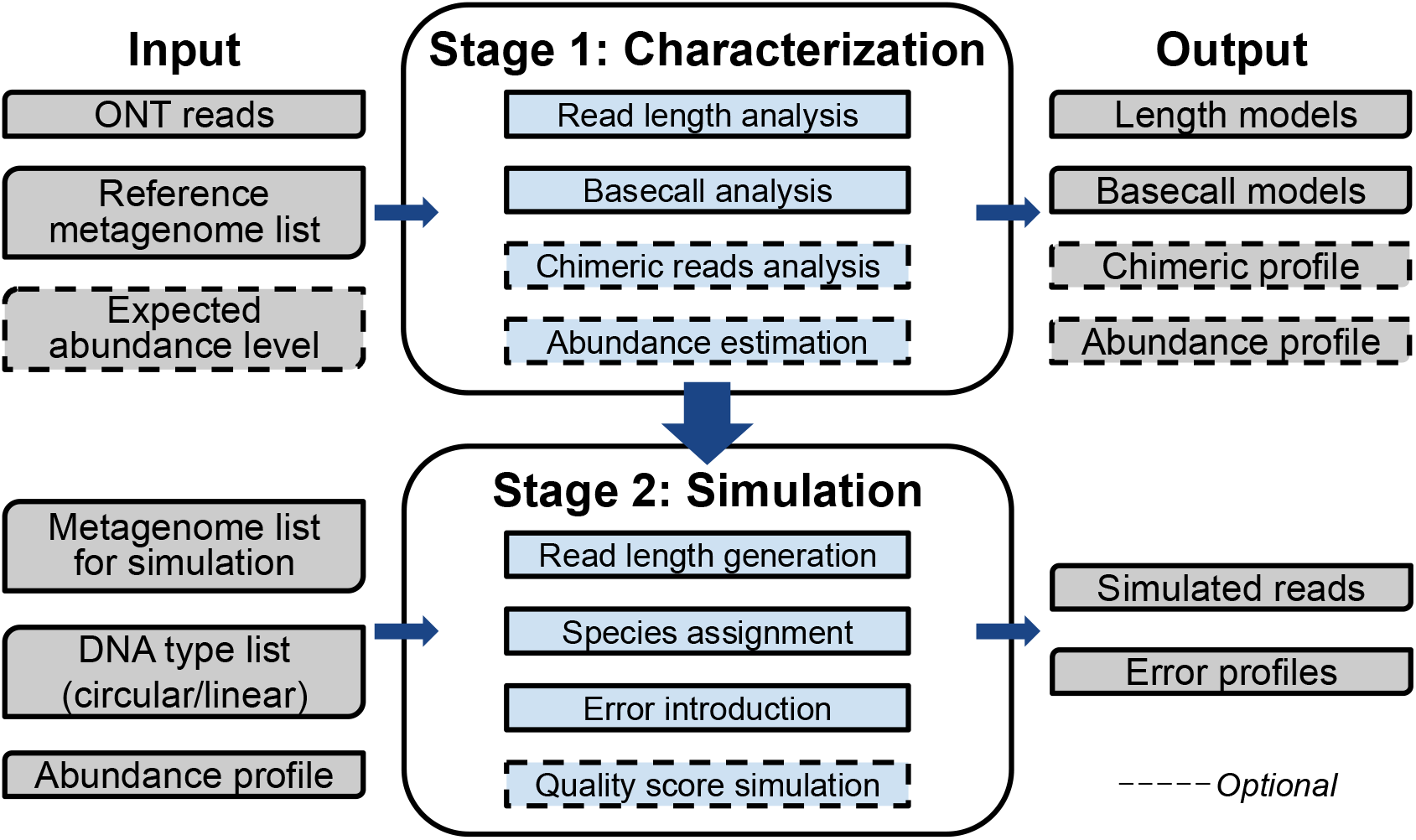
Meta-NanoSim workflow. Meta-NanoSim consists of two stages: characterization stage and simulation stage. In the characterization stage, given a training dataset and reference metagenome, Meta-NanoSim builds models for the read length distributions and basecall events. It optionally profiles chimeric read artifacts and quantifies an abundance profile. It can also calculate the deviation between expected and estimated abundance levels. In the simulation stage, Meta-NanoSim takes four inputs: 1) The genome list for simulation (local or ftp path), 2) DNA type list, 3) abundance profile for simulation, and 4) the models generated from the characterization stage. Meta-NanoSim outputs simulated reads and error profiles as ground truth.

To simulate a metagenomic dataset, besides the pre-trained model from the characterization stage, a list of reference genomes of users’ choice is required as input to use for simulation, together with their abundance levels and DNA topology (i.e. linear or circular). The tool can also stream reference genome sequences from either RefSeq (30) or Ensembl (31) automatically without requiring extra disk storage, which facilitates large microbial community simulations. Since microbial sequencing projects are often carried out in a multi-sample or multi-replicate fashion, Meta-Nanosim is designed to simulate multiple samples in one batch with user-defined abundance level profiles as input.

### Chimeric read characterization and simulation

Chimeric reads are also called split reads because one read is split into two or more sub-alignments that are aligned to distinct regions of the genome / metagenome. They may be created due to sequencing artifacts or structural variants when the reference metagenome is not comprehensive. Previous studies reported that chimeric reads represent a non-negligible fraction of ONT sequencing datasets ranging from 1.7% to 8.17% with different seuqncing kit and identificaiton thresholds (17,20,22,23). In the metagenome datasets used in our study, after ruling out structural variants, we have identified a similar fraction of reads in this category: 2.17% (75,628 reads) in the *Even* dataset and 1.67% (68,444 reads) in the *Log* datasets. These reads are free of known adapters, so their presence may impact downstream analyses, such as assembly, taxonomy binning, and quantification, even after adapter trimming. When aligned to their respective reference genome sequence(s), ONT reads may contain unaligned or soft-clipped regions. In our tests, the chimeric read detection feature of Meta-NanoSim significantly reduced the length of these unaligned regions, which explained why some of the reads have over 1 kbp long unaligned portions (Fig. 2A). As seen in Fig. 2B, the length distributions of the gaps between split alignments follow multi-modal distributions. Meta-NanoSim uses kernel density estimation to model them, with results exhibiting strong similarity between the length distributions of simulated and experimental sequences. We also noticed that the number of segments each read contains can be described as a geometric distribution and the mean probability can be learnt from experimental data (Fig. 2C). On average, each read contains 1.03 segments for both data sets under study.

**Fig. 2.**
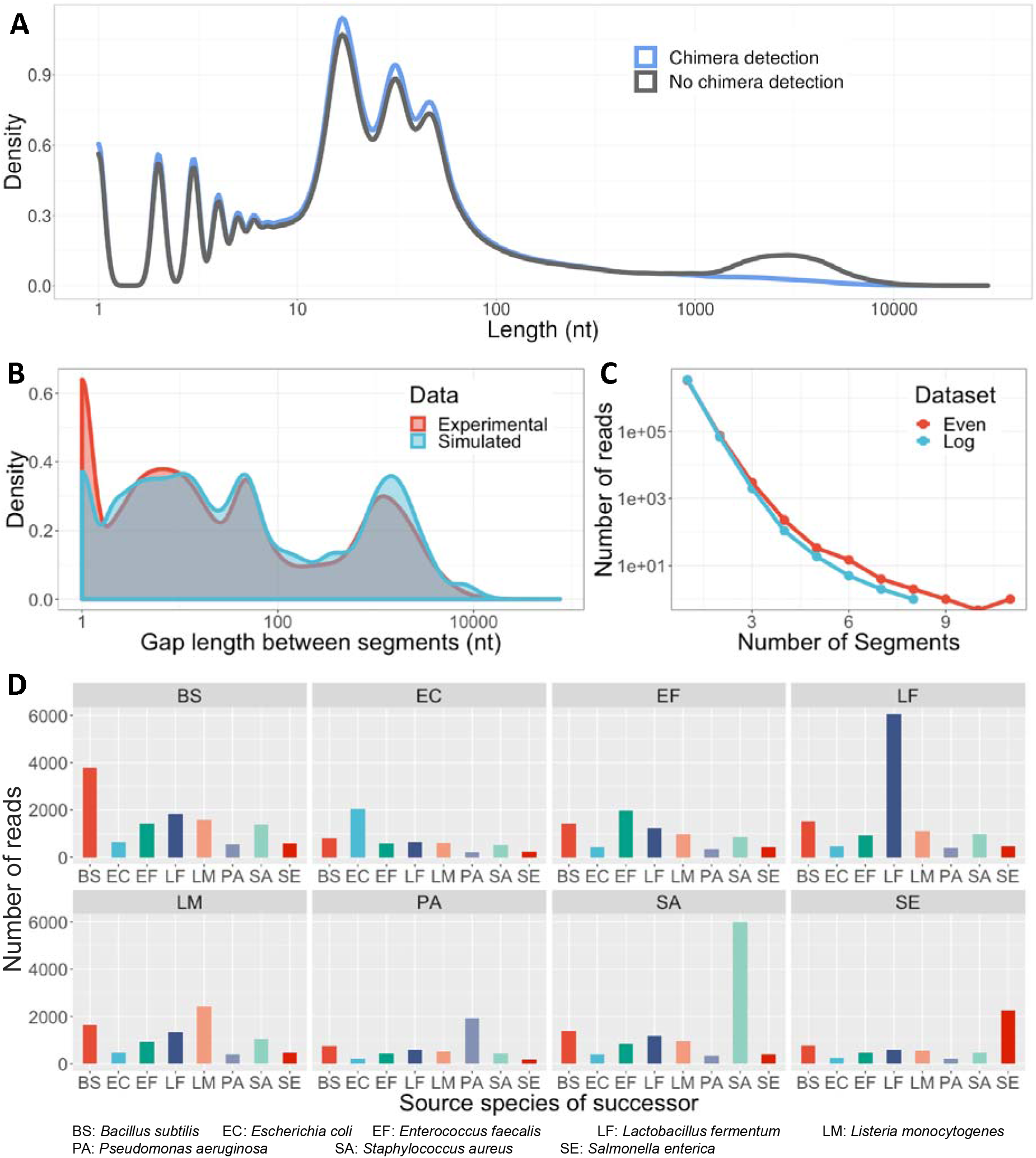
Chimeric reads detection and simulation. **A.** The length distribution of the unaligned regions of reads with or without chimeric reads detection for the *Log* dataset. **B.** The performance of gap length simulation for the *Log* dataset. **C.** The number of segments each read contains for the *Even* and *Log* datasets. **D.** All segments in chimeric reads in the *Even* dataset are converted into overlapping pairs. Each facet represents one source species of the first segment and the *x*-axis represents the source species of the second segment. Each facet shows the probability of the second segment given the source species of the first one. *Cryptococcus neoformans* and *Saccharomyces cerevisiae* are excluded here due to their low abundances.

Based on the source species to which each split alignment belongs, chimeric reads can be classified as “intra-species-chimeric” or “inter-species-chimeric”. It is observed that the source species of the first segment is affected by the abundance level, while subsequent segment is more likely to be influenced by the identity of the previous species (Fig. 2D). We postulate that this is because DNA molecules of the same species are more likely to gather near the nanopore than being homogeneously dispersed in the buffer. Regardless of the actual cause, this phenomenon can be approximated as a simplified hidden Markov model with a generalized emission probability, which is defined as shrinkage rate *s* here (Methods). To our calculation, s is equal to 0.77 for the *Even* dataset and 0.73 for the *Log* dataset, suggesting that its value is stable across each dataset.

### Abundance estimation

Existing abundance estimation methods generally quantify the number of mapped reads or *k*-mers, under the presumption that all reads have equal lengths. However, since the ONT read length varies several folds of magnitude, it is likely that the mean read length for each speces is different. When all species are equally and deeply sequenced, according to central limit theorum, the standard deviation of mean lengths would scale with 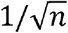, where n is the number of species. In reality, low-abundant species may have a even higher standard deviation because there are fewer sequences representing it. We observe the mean read lengths of uniquely aligned reads in the *Log* datasets to range between 3-7kb, neccesitating base-level instead of read-level quantification algorithms (Fig. S2). One key challenge that confounds short-read metagenomic analysis is ambiguously aligned reads. In ONT datasets, however, most reads are long enough to span inter- and intra-species homologous regions, and the chimeric reads detection feature can resolve the estimation for reads that have multiple sub-alignments and for reads that span across the start site of a circular genome. For the remaining small fraction of multi-aligned reads between closely-related species, the estimation for them can be optimized using the EM algorithm (Methods).

To benchmark, we compared the performance of four abundance estimation methods: Meta-NanoSim estimation with chimeric reads detection, Salmon quantification with `--meta` option (Salmon) (32), the base-level estimation reported in the paper that released the dataset (denoted as “Data Note” from hereon) (8), and MetaMaps. For Meta-NanoSim estimation, we performed an ablation study that removes key components of the algorithm step by step, including estimation on read-level with chimeric reads detection (Meta-NanoSim CR) or with EM algorithm (Meta-NanoSim ER), estimation on base-level (Meta-NanoSim B), base-level with chimeric reads detection (Meta-NanoSim CB), base-level with EM algorithm (Meta-NanoSim EB), and base-level with chimeric reads detection fine-tuned by EM algorithm (Meta-NanoSim ECB). All compared methods, except for MetaMaps, are computed based on Minimap2 alignments. We compared the estimated abundances to the expected ones provided by the manufacturer and computed all the metrics: R-squared, standard deviation, and percent error.

In general, all base-level quantification methods performed better than read-level quantification methods (Salmon, MetaMaps, Meta-NanoSim CR and ER), and Meta-NanoSim base-level estimations have the highest correlation with the expected abundances (Table 1, Fig. S3). For the *Even* dataset, all four Meta-NanoSim base-level methods performed similarly; the stand-alone base-level quantification has the highest R-squared value for the *Even* dataset, while the chimeric reads detection helped reduce the percent error, mainly for low-abundance species *Cryptococcus neoformans* (Fig. S3). MetaMaps, as a read-level quantification method designed specifically for ONT metagenomic data, although ranked highest among this category, showed a big discrepancy compared to base-level methods with a near-doubled percent error. For the *Log* dataset, Meta-NanoSim base-level estimations also had similar R-squared values, higher than other methods. Since the metrics are close to each other for the *Log* dataset and the performance on low-abundance species may be overshadowed by high-abundance species, we also computed the coefficient of correlation and error between log-transformed estimated and expected abundances. After log-transformation, Salmon, Meta-NanoSim ECB, and Meta-NanoSim EB showed similar performance with Salmon having the highest correlation and Meta-NanoSim ECB having the lowest percent error. The metrics for Meta-NanoSim estimation without EM, on the other hand, dropped significanly due to difficulties in differentiating multi-mapped reads for low-abundance species. When estimating the abundance levels for the *Log* dataset, Minimap2 incorrectly assigned 18,212 reads to the *E. faecalis* genome as primary alignments, when they can also be aligned to an inter-species homologous region in the *L. monocytogenes* genome. In fact, *E. faecalis* is a low-abundance species in the *Log* dataset with only 33 unique alignments. Therefore, the methods with EM algorithm resolved the multi-aligned reads problem, indicating that EM can be advantageous for datasets consisting of similar genomes but with large variances in abundance levels.

**Table 1.**
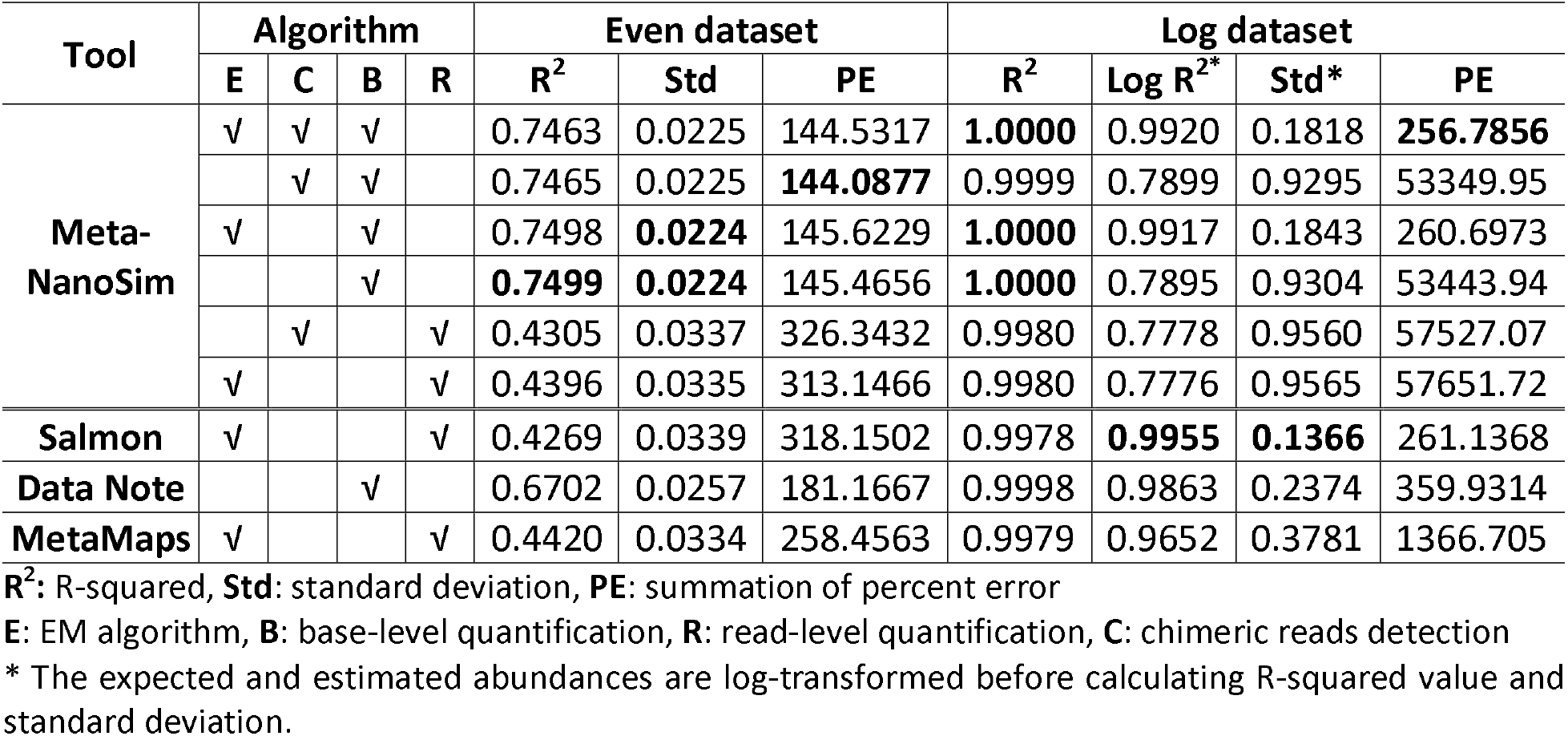
Statistical analysis of the abundance estimation results compared to expected abundances.

To recapitulate our findings, we also chose another logrighmically distributed mock microbial community, denoted as the *Adp* dataset (33) (Methods), and repeated the quantifications with Meta-NanoSim base-level methods, Salmon, and MetaMaps (Table S1). Similarly, all four Meta-NanoSim methods performed similar to each other with the highest correlation to the expected values. MetaMaps quantification, although had the lowest percent error among all compared methods, showed a much lower correlation in terms of R-squared and standard deviation. Taken all together, Meta-NanoSim base-level quantification after chimeric read detection and fine-tuned by EM algorithm balanced correlation and percent error well, making it preferrable for naturally occurring microbial communities with various abundances.

Because of the deviation between expected and estimated abundance levels, we introduce a feature that can simulate this observation. With user-provided deviation range, Meta-NanoSim first varies the abundance levels to desired fluctuation, and then starts the simulation process (Methods). We compared the deviation between expected abundances and experimental data, and the simulation results of NanoSim and CAMISIM (Fig. 3). The distribution of abundance deviations of experimental data and Meta-NanoSim simulated reads are statistically the same (Kolmogorov-Smirnov test *p*-value = 0.79), while the one of CAMISIM reads is pronouced as it does not provide this feature.

**Fig. 3.**
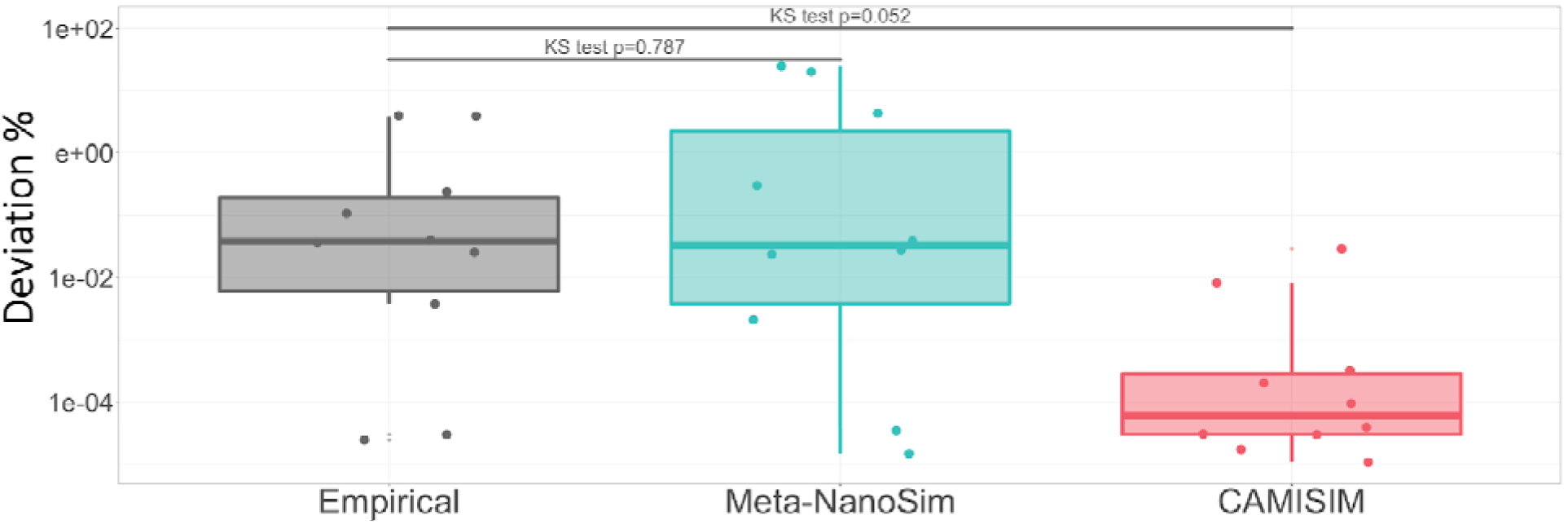
The simulation of abundance level deviations. In this plot, each dot represents microbial genome, and the y-axis represents the deviation in percentage between the expected values and experimental / simulated values.

### Comparison between simulated and experimental datasets

To demonstrate the performance of Meta-NanoSim, we trained it with the *Log* dataset and compared the simulated datasets against the result of CAMISIM. With eight processors, simulation of one million reads took under 20 minutes (or under 160 CPU-minutes) for Meta-NanoSim, while CAMISIM required more than six hours to complete.

The read lengths of simulated datasets from Meta-NanoSim follow the empirical length distribution closely, with a median read length peak at 4,040 nt (3,994 nt for empirical reads) (Fig. 4A). In contrast, the lengths of CAMISIM-simulated reads deviate far from those of the empirical data. Because CAMISIM is only compatible with an old version of NanoSim and uses hard coded pre-trained models learnt from a genomic sequencing data, it was not possible to test it coupled with Meta-NanoSim. Moreover, the length distribution of unaligned parts on Meta-NanoSim simulated reads captures the patterns in empirical reads well, with multiple peaks below 100 nt. In contrast, the lengths of unaligned part in CAMISIM reads are inflated as it does not detect nor simulate chimeric reads. Both Meta-NanoSim and CAMISIM mimic the mismatch and deletion events well when compared to the empirical dataset (Fig. 4B), which demonstrates the robustness of NanoSim mixture statistical models. However, Meta-NanoSim simulates insertion and match events better than CAMISIM, also due to the change of model.

**Fig. 4.**
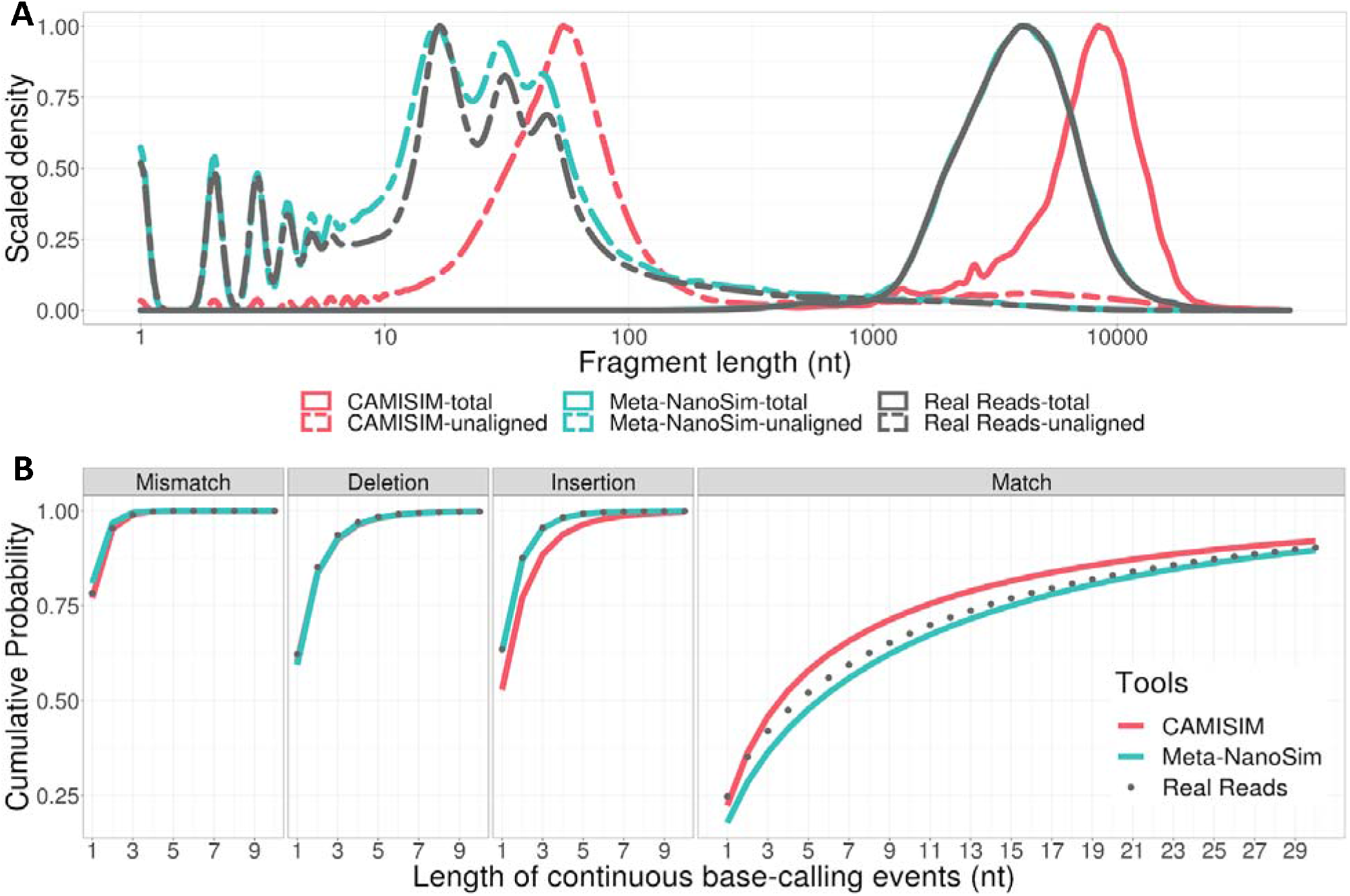
Performance of Meta-NanoSim and CAMISIM in simulating one million reads from the *Log* dataset. **A.** Comparison of read length distributions in the empirical vs. simulated reads. Unaligned length represents the length of unaligned part of each aligned read. **B.** Cumulative probability function of the lengths of matches/errors in empirical and simulated reads.

In addition, we challenged Meta-NanoSim with two more simulation tasks. First, we simulated two samples at the same time with the pre-trained model from the *Log* dataset. Each sample contained one million reads from the seven species from the *Adp* dataset with different abundance levels. Meta-NanoSim simulation finished within 51 min with eight processors. Although the metagenome to be simulated is completely different from the training one, simulated reads exhibited similar read features as the experimental data, and the abundance levels are 100% in accordance with the expected values (Fig. S4). Next, we randomly picked a saliva sample from the Human Microbiome Project (HMP) and tried to simulate ONT reads using the same microbial composition (34). With 125 different bacteria straines, it took Meta-NanoSim less than three hours to simulate 10 million reads, including the time used for streaming reference genomes from RefSeq server.

### Benchmarking on assembly tasks

Meta-NanoSim simulated reads can be used to benchmark and assess the performance ofbioinformatics algorithms. To demonstrate an application of Meta-NanoSim, we simulated four sets of data with 1, 2, 4, 10 million reads to assess the correctness, scalability and robustness of metaFlye, a long-read metagenomic assembler. We used the models learnt from the Log dataset and the abundance levels of the 10 species are shown in Fig. 5. With 128 threads, runtimes ranged from one hour to seven hours (Table S2). The maximum resident set size for 10 million reads dataset was 212 GB, but intermediate files occupied over 10 TB of disk space during consensus-building stage. We also tried to assemble a larger dataset of 20 million reads, however the assembly failed after 30 days with an out-of-memory error on a 1 TB RAM server. Before completing, the maximum resident set size was 1.75 TB, and the most time-consuming stage was the graph simplifying stage (two weeks).

**Fig. 5.**
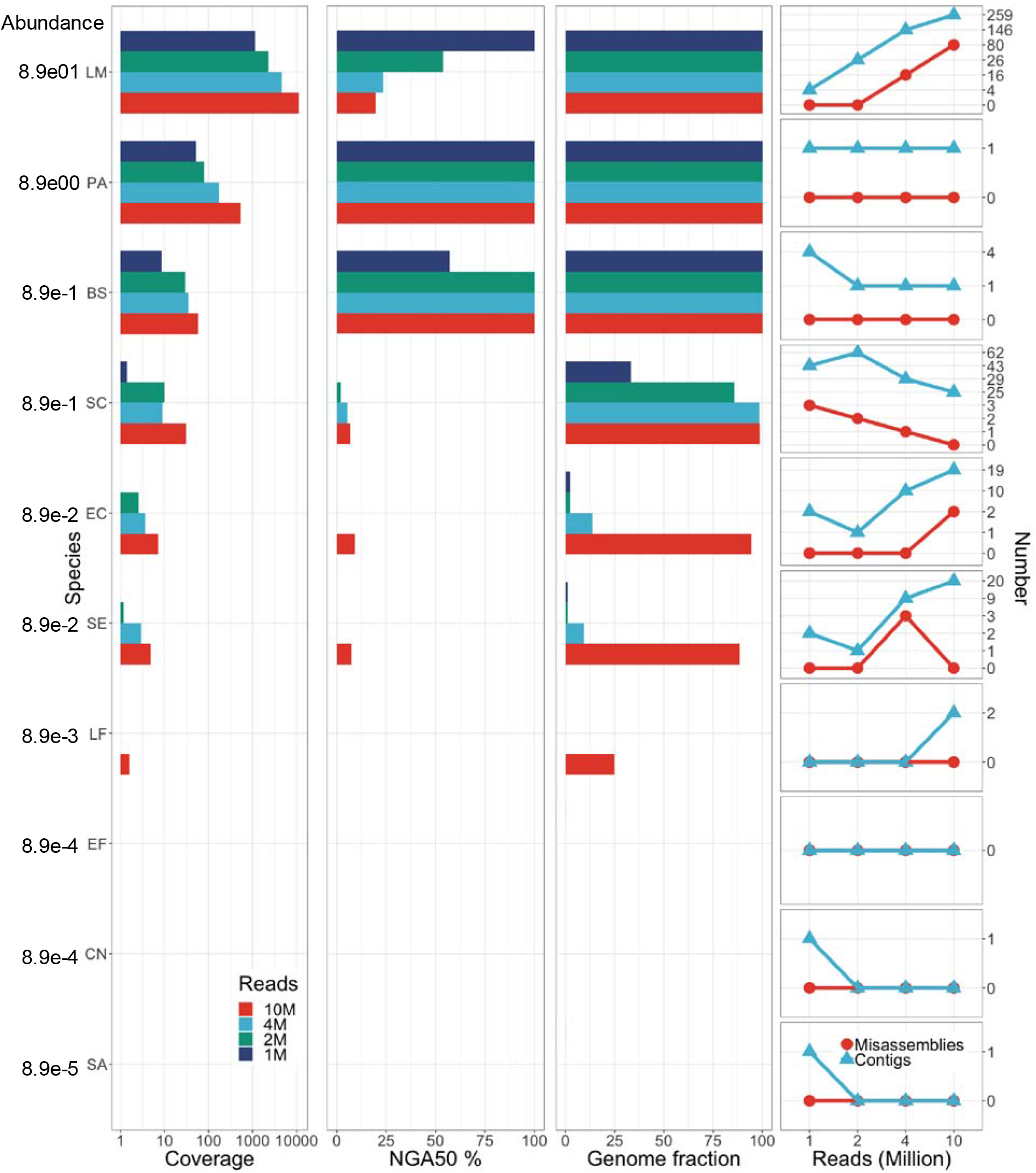
metaFlye assemblies with four sets of simulated datasets. Four sets of simulated datasets include 1, 2, 4 and 10 million reads, respectively. The abundance is the expected abundance level during simulation. Coverage panel is the average read depth including plasmids. NGA50 % represents the NGA50 length divided by the reference genome size. Genome fraction is a proportion between the assembled sequences and each corresponding genome. The right-most panel shows the number of misassemblies and assembled contigs as the number of simulated reads increases. BS: *Bacillus subtilis*, CN: *Cryptococcus neoformans*, EC: *Escherichia coli*, EF: *Enterococcus faecalis*, LF: *Lactobacillus fermentum*, LM: *Listeria monocytogenes*, PA: *Pseudomonas aeruginosa*, SA: *Staphylococcus aureus*, SC: *Saccharomyces cerevisiae*, SE: *Salmonella enterica*.

In total, MetaFlye reconstructed 27.81 Mbp sequences, adding up to 43.72% of the total reference genome for the 4M reads simulated dataset. These metrics are similar to the reported assemblies using the original training dataset with 3.48 M reads (28.20 Mbp assembled length that covered 46.00% of the reference metagenome) (29). Generally speaking, the average genome coverage is positively correlated with the number of sequencing reads and abundance levels, and accordingly the genome reconstruction fraction and NGA50 length are positively correlated with the coverage (Fig. 5). As expected, sub-1x coverage genomes have very poor reconstructions. Between 1x and 10x, the positive correlation is mirrored in multiple species, including *B. subtilis, S. cerevisiae, E. coli* and *S. enterica*. When the coverage reaches 10x, metaFlye is able to reconstruct the genome to nearly 100% (*S. cerevisiae* in the 4M dataset, Fig. 5). When the coverage reaches 30x, the NGA50 length can cover the whole genome size (*B. subtilis* in the 2M dataset). Similarly, the contigs reconstructed for these two species decrease as the number of reads increase, showing how increasing sequencing depth can help assembling into one contig for *B. subtilis* and nearly one contig per chromosome for *S. cerevisiae*. In contrast, when the coverage is not high enough, the assemblies may be fragmented or even mis-assembled in case of *E. coli* and *S. enterica*. However, a higher coverage does not necessarily lead to a better assembly quality. The reconstruction of *L. monocytogenes* deteriorates with more reads when the fold coverage exceeds 1000x. Although the genome fraction remains 100%, the NGA50 is only half or less of the genome size for the 2M, 4M, and 10M datasets. The drop in NGA50 length can be explained by the increasing number of reconstructed contigs and mis-assemblies (Fig. 5). With one million reads, there are only four contigs that can be mapped to the *L. monocytogenes* genome, and there are no mis-assemblies detected. However, we believe that the higher sequencing depth above 1000X has led to many more misassemblies, which lowered the assembly’s NGA50.

## DISCUSSION

The applications of nanopore sequencing on metagenomic projects are rapidly expanding, motivating the development of metagenomic analysis tools tailored for this specific data type. In this work, we bring two main contributions to ONT metagenomic analysis tasks: (1) introduction of a new base-level quantification method for metagenomic abundance estimation; and (2) an upgrade of NanoSim for metagenomic characterization and simulation.

Reference-based metagenomic abundance estimation is key to investigating the microbial composition of an enviroment, and this has been a recurring topic for all emerging sequencing technologies. The long read length of ONT reads provides an opportunity to resolve homologous regions between species or even strains, but other complications arise due to the high error rate and non-uniform read lengths. Existing methods generally assume uniform read lengths and therefore they only need to count the number of mapped reads or *k*-mers. However, it is unreasonable to treat, say, a 100 bp long read and a 1000 bp long read the same when calculating their contributions to the genome abundance. Our results echoed that it is necessary to quantify microbial abundances on a base-level rather than read-level to leverage this data type. In addition to the higher error rate, a small yet substantial fraction of ONT reads are chimeras, which may obscure the accuracy of estimates. The chimeric read detection feature in Meta-NanoSim works by searching for best compatible alignments and helps reducing the percent error in abundance estimation. For multi-aligned segments whose source species cannot be differentiated, we adopted an EM algorithm to optimize their proportional contributions to each potential source species. Our benchmarking results demonstrate that the combination of these three components can achieve the best correlation with expected abundance. We note that Meta-NanoSim quantification performs better when the abundance levels are more uniform or when low-abundance microbes do not share large homologous regions with high-abundance microbes. Depending on wheather the user wishes to achieve a higher correlation or lower percent error, they can choose to disable or enable chimeric reads detection, respectively. Although our work is limited to reference-based quantification, we expect it to elucidate the design of reference-free methods to have a broader application.

Built on top of the abundance estimation, Meta-NanoSim is able to simulate datasets with desired abundance profiles. It can also recapitulate the abundance level deviation from expected values, which is especially useful for designing sequencing projects. When the rough abundance of a microbial community is known, it is essential to know how deep the sequencing needs to be so that each species can be covered by a sufficient number of sequencing reads. However, when the sequenced abundance differs from the expected value, simulated data with abundance variations true to the platform can inform the relationship between sequencing depth and abundance levels.

The general workflow of characterization and simulation of Meta-NanoSim follows the same paradigm of the previous versions of NanoSim. The chimeric read detection in characterization stage provides a means to profile all chimeric reads in a library regardless of its root cause. When the reference metagenome is inclusive, the chimeric reads are likely introduced by library preparation and sequencing artifacts; while in reality, since the detection relies on alignment, some chimeric reads may also be attributed to structural variants when the source genome is not present in the reference. In this case, the output of the characterization stage can be used to further investigate such events with specifically designed statistically models and algorithms. Three main new features for simulation added to Meta-NanoSim are chimeric read simulation, streaming reference genome for online server, and the simulation of a metagenome composed of a mixture of both linear and circular genomes. As chimeric reads may interfere with downstream analyses, simulated datasets with these artifacts are needed for more accurate performance assessment. Characterizing this feature and introducing it to simulated reads will also diversify error types in the reads, helping to improve the robustness of related algorithms. Reference genome streaming is uniquely advantageous when simulating a large metagenome with hundreds of species. It eliminates the reference genome downloading step for users and saves disk space while keeping the runtime reasonable. Similarly, since metagenomes are naturally composed of both linear and circular genome topologies, having a simulated dataset supporting this important characteristic will add credibility to benchmarking results and better forecast performance with experimental data.

The benchmarking on metagenomic assembler showcased that Meta-NanoSim can facilitate relevant tool development as well as guide sequencing projects. The resulting assembly quality of Meta-NanoSim simulated reads is comparable to that of the experimental data with similar coverage. Although publicly available mock community sequencing data provide a more realistic training and test set, simulated data provide a ground truth and has no limit in size, making them perfect for testing the accuracy and scalability of algorithms. Through the use of simulated datasets, we demonstrated that metaFlye assembler performs best when the species coverage is between 10 and 1000-fold. To ensure a successful assembly of low abundance species, it is suggested to calculate the number of reads needed given an estimated abundance first to ensure just enough coverage without wasting resources. For example, it takes 10 million reads to achieve 10-fold coverage for a 0.1% abundant species with a genome size of 5Mb. When assembling real microbial communities with highly variable abundance levels, we recommend multiple rounds of assembly with different sample sizes to achieve the best performance for both high- and low-abundance microbes. In addition, developers may analyze in depth the mis-assemblies and errors in assembled contigs with the ground truth provided by Meta-NanoSim to improve their algorithms. The effect of chimeric reads, as a common source of mis-assemblies, can be easily evaluated with simulated reads. Meta-NanoSim also has a perfect read simulation feature, which would allow users to learn the relationship between assembly and coverage without interference from read base errors. Furthermore, Meta-NanoSim simulated datasets can serve as suitable training and testing datasets for a wide range of algorithms, including read error correction, taxonomy binning, and compositional estimation.

## CONCLUSIONS

Meta-NanoSim is an ONT metagenomic simulator that simulates complex microbial community with read features true to the platform. Given a training dataset, Meta-NanoSim generates read length distributions, error profiles, and alignment ratio models by default. It also detects chimeric reads and quantifies species abundance levels on demand. Meta-NanoSim is aimed to capture the platform-specific features and can be adopted to profile datasets from any sequencing chemistry and basecallers. The performance of metagenomic quantification of Meta-NanoSim surpasses the performance of the current state-of-the-art. Chimeric read detection improves the read length modelling and helps to reproduce such feature in simulated reads to challenge metagenomic assemblers, taxonomy binners, and abundance quantification tools. For the simulation part, Meta-NanoSim is the first ONT metagenomic read simulator that can simulate chimeric reads and abundance levels at base-level. It also supports multiprocessing and streamed reference genomes from online servers to speed up simulations when hundreds or thousands of genomes are to be simulated in a microbial community. By comparing simulated reads with empirical datasets, we show that Meta-NanoSim preserves some key characteristics of ONT metagenomic reads well. Further, our metagenomic assembly benchmarks demonstrate the usability and utility of Meta-NanoSim and we expect Meta-NanoSim to have broad utility in helping develop, test and improve upon such applications.

## METHODS

### Datasets

For training and testing meta-NanoSim, we used two publicly available ONT datasets generated on the GridION platform (8). ERR3152364 was sequenced from the Zymo CS Even ZRC190633 community, and ERR3152366 was sequenced from the Zymo CSII Log ZRC190842 community. Both communities contain eight bacteria species and two fungi species; the abundance levels are 12% for each bacterium and 2% for each fungus for the *Even* dataset; and the abundance levels for the *Log* dataset are shown in Fig. 3A. Reads were basecalled using Guppy v2.2.2 GPU basecaller with flip-flop configuration, and adapters were trimmed using Porechop v0.2.4. Reads were also split when internal adapters were found to eliminate false chimeric reads. Additionally, we used another publicly available ONT dataset, the *Adp* dataset, from project PRJEB44844. This dataset is composed of three libraries with different mean sequence length: ERR5897838, ERR5903399, and ERR5909878. Reads from this dataset were basecalled during the experiment on the GridION using the MinKNOW software. The abundance levels are 50% *A. xylosoxidans*, 25% *M. morganii*, 12% *L. richardii*, 6% *P. aeruginosa*, 4% *M. wisconsensis*, 2% *P. vulgaris* and 1% *S. dysgalactiae*.

The experimental dataset for naturally occurring microbial community is from the HMP project, human saliva sample SRS019120. The abundance table of the sample was downloaded from https://hmpdacc.org/hmp/HMSCP/. In the original file, there were 176 bacteria strains, and after removing the ones without reference genome in RefSeq, we compiled a list of 125 strains with their RefSeq reference genome ftp site. We computed the abundance levels based on the size of the reference genome and the estimated depth in the abundance file, and fed it into Meta-NanoSim for simulation. The simulated 10 million reads are uploaded to Zenodo for academic use: https://doi.org/10.5281/zenodo.5712441.

### Chimeric reads detection and simulation

During alignment, one ONT read may have multiple sub-alignments to different *loci* on the reference genome / metagenome. When the query and reference coordinates of two sub-alignments do not overlap, we define them as compatible alignment of each other. Finding the best compatible alignment set problem is akin to finding the largest compatible interval set in computer science. For each read, we exhaustively search for all compatible alignments for each sub-alignment to generate a list of compatible alignment sets (Fig. S1). We then select the best element from the list for downstream analysis, based on alignment quality and total aligned length. If, for a given read, the best element contains two or more compatible alignments, the read is considered as chimeric and its aligned length, gap length, and source species (in metagenome mode) are modelled for simulation. One exception here is reads that span the start position of a circular genome. These reads are detected but not considered chimeric. The two sub-alignments are cancatenated as one long alignment to count towards the aligned length and source species.

To determine the source species for each segment in chimeric reads, we build a simplified hidden Markov model where the start probability is the input abundance, the emission probability represents which species the next segment is coming from given the previsous one, and the transitional probability of species is the change of abundance in the underlying Markov chain. Since species to be simulated, namely the states in a Markov model, may be different between the training and simulation metagenome, we generalize the emission probability as a single value called shrinkage rate *s* (0 < *s* <= 1). This parameter describes the reduction of abundances (probabilities) of other species, while maintaining the relative abundances among them. Assuming the input abundance is {*p*_*A*_, *p*_*B*_, *p*_*C*_, ..., *p*_*N*_} for n species, when the first segment comes from species A, the transitional probabilities for the other species would become {*s×p*_*B*_, *s×p*_*C*_ ... *s×p*_*N*_} and the transitional probability for A would be inflated as 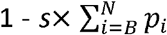. To learn *s*, all segments in chimeric reads are divided into overlapping pairs, and the probability for the source species of the second segment being different from the first one is recorded. In this way, we can calculate the reduction of abundance for every species. The average reduction is the shrinkage rate, and the inflated abundance for being from the same species can be inferred as well. The shrinkage rate can also be adjusted by the user, to 1 for example, if one assumes all DNA molecules are homogeneously suspended in the buffer.

To summarise the simulation of chimeric reads, Meta-NanoSim first determines the number of segments to be simulated based on the Geometric distribution. If the number of segments is greater than or equal to two, it is a chimeric read. Then, Meta-NanoSim generates the lengths of each segment and gap between them using kernel density estimation learnt from empirical reads. The source species of the first segment is randomly picked based on the input abundance level. Starting from the second segment, the abundance levels are re-computed based on the previous species and *s*. The source species is determined one after another, and then sequences are extracted, mutated with purposely introduced errors, and assembled in the same process as non-chimeric reads.

### Abundance estimation

Meta-NanoSim offers two estimation methods, one with the chimeric reads detection and one without. When chimeric reads detection is enabled, all subalignments are used for computing; when it is disabled, only the primary alignments are used. Meta-NanoSim records the aligned bases for each sub-alignment towards their source genome, and then uses EM algorithm (Supplementary Methods) to assign multi-aligned segments proportionally to their putative source genomes iteratively.

### Abundance deviation simulation

Meta-NanoSim offers a means to simulate the abundance deviation with user-defined lower and upper deviation boundaries. We noticed a weak positive correlation between genome size and abundance deviation in our analysis. During simulation we first randomly draw a list of relative error between the deviation boundaries. Next we assign these errors to each genome based on their sizes, namely larger errors are assigned to larger genomes and smaller errors are assigned to smaller genomes. After modifying with errors, the abundances are renormalized to have a total abundance of 100%.

### Abundance estimation benchmarking

For the Data Note estimation, we used the data provided in the paper where the dataset was first reported (8). For all other three methods, we first mapped the reads to the ZymoBIOMICS reference genome version 2 with Minimap 2.17-r941. To run Salmon, we used the `--meta` option and `--noErrorModel` to quantify the abundance for both datasets as suggested in the manual.

### Simulation and benchmarking

We trained Meta-NanoSim on the mock community datasets with two options, `--chimeric` and `--quantification (src/read_analysis.py metagenome -i input.fa -gl metagenome_list_for_training -t 12 -q -c)`, and then simulated one million reads with `--chimeric (src/simulator.py metagenome -gl metagenome_list_for_simulation -a abundance_for_simulation.tsv - dl dna_type_list.tsv -c models/training -t 12 -c)`. The abundance levels simulated are the same as the *Log* dataset and *Adp* dataset, when respective metagenome were used for simulation. CAMISIM uses NanoSim as the engine for ONT read simulation. The pre-trained profile is hardcoded in CAMISIM, which is trained on an *E. coli* dataset with NanoSim version one. To run CAMSIM, we had to choose a most updated compatible version of NanoSim (V2.0.0) that reads the specific format of profiles. Then we mapped the simulated datasets to the reference genomes with Minimap 2.17-r941 to calculate the read length distributions and error distributions, and compared the result with the ones of the experimental data.

### Assembly benchmarking

The assembly benchmarking was performed on a high-performance computing server with 128 CPUs and 1 TB memory. We simulated four sets of data with models trained on the *Log* dataset with default settings and each dataset contains 1, 2, 4, 10 million reads. Then we ran metaFlye 2.8.1-b1676 with options `--meta, --plasmids, --threads 128` and `-g 70m`. Next we ran MetaQUAST v5.1.0rc1 with default settings to evaluate the quality of assemblies.

## LIST OF ABBREVIATIONS

bp: basepairs
EM: Expectation-Maximization
GB: gigabytes
GPU: graphics processing unit
M: million
NGA50: length of the shortest alignment block for which longer or equal length alignment blocks cover 50% of the reference genome size
nt: nucleotides
ONT: Oxford Nanopore Technologies
TB: terabytes

## DECLARATIONS

### Ethics approval and consent to participate

Not applicable

### Consent for publication

Not applicable

### Availability of data and material

Meta-NanoSim is implemented in Python within the NanoSim suite. The source code and pre-trained models used in this study are available on Github: https://github.com/bcgsc/NanoSim. NanoSim version 3.0.2 is used for this work. Meta-NanoSim is platform independent, and is released under the GNU GPL license.

### Authors’ contributions

IB and CY conceived and designed the study. CY designed and implemented the software with the help of TL, SH, and KMN. KMN and SH provided additional help with the software maintainence. CY drafted the manuscript, and all authors were involved in its revision. All authors read and approved the final manuscript.

## Acknowledgements

Not Applicable

## REFERENCE

1. Handelsman J. Metagenomics: Application of Genomics to Uncultured Microorganisms. Microbiol Mol Biol Rev. 2004;

2. Chen K, Pachter L. Bioinformatics for whole-genome shotgun sequencing of microbial communities. PLoS Computational Biology. 2005.

3. Schulz F, Alteio L, Goudeau D, Ryan EM, Yu FB, Malmstrom RR, et al. Hidden diversity of soil giant viruses. Nat Commun. 2018;

4. Guthrie L, Gupta S, Daily J, Kelly L. Human microbiome signatures of differential colorectal cancer drug metabolism. npj Biofilms Microbiomes. 2017;

5. Wirbel J, Pyl PT, Kartal E, Zych K, Kashani A, Milanese A, et al. Meta-analysis of fecal metagenomes reveals global microbial signatures that are specific for colorectal cancer. Nat Med. 2019;

6. Quince C, Walker AW, Simpson JT, Loman NJ, Segata N. Shotgun metagenomics, from sampling to analysis. Nature Biotechnology. 2017.

7. Brown BL, Watson M, Minot SS, Rivera MC, Franklin RB. MinIONTM nanopore sequencing of environmental metagenomes: A synthetic approach. Gigascience. 2017;

8. Nicholls SM, Quick JC, Tang S, Loman NJ. Ultra-deep, long-read nanopore sequencing of mock microbial community standards. Gigascience. 2019;

9. Fu S, Wang A, Au KF. A comparative evaluation of hybrid error correction methods for error-prone long reads. Genome Biol. 2019;

10. Payne A, Holmes N, Rakyan V, Loose M. Bulkvis: A graphical viewer for Oxford nanopore bulk FAST5 files. Bioinformatics. 2019;

11. Charalampous T, Kay GL, Richardson H, Aydin A, Baldan R, Jeanes C, et al. Nanopore metagenomics enables rapid clinical diagnosis of bacterial lower respiratory infection. Nat Biotechnol. 2019;

12. Kafetzopoulou LE, Pullan ST, Lemey P, Suchard MA, Ehichioya DU, Pahlmann M, et al. Metagenomic sequencing at the epicenter of the Nigeria 2018 Lassa fever outbreak. Science (80- ). 2019;

13. Chan JFW, Yuan S, Kok KH, To KKW, Chu H, Yang J, et al. A familial cluster of pneumonia associated with the 2019 novel coronavirus indicating person-to-person transmission: a study of a family cluster. Lancet. 2020;

14. Greninger AL, Naccache SN, Federman S, Yu G, Mbala P, Bres V, et al. Rapid metagenomic identification of viral pathogens in clinical samples by real-time nanopore sequencing analysis. Genome Med. 2015;

15. Yang C, Chu J, Warren RL, Birol I. NanoSim: Nanopore sequence read simulator based on statistical characterization. Vol. 6, GigaScience. 2017.

16. Hafezqorani S, Yang C, Lo T, Nip KM, Warren RL, Birol I. Trans-NanoSim characterizes and simulates nanopore RNA-sequencing data. Gigascience. 2020;

17. Buck D, Weirather JL, de Cesare M, Wang Y, Piazza P, Sebastiano V, et al. Comprehensive comparison of Pacific Biosciences and Oxford Nanopore Technologies and their applications to transcriptome analysis. F1000Research. 2017;

18. Wood DE, Salzberg SL. Kraken: Ultrafast metagenomic sequence classification using exact alignments. Genome Biol. 2014;

19. Lu J, Breitwieser FP, Thielen P, Salzberg SL. Bracken: Estimating species abundance in metagenomics data. PeerJ Comput Sci. 2017;

20. White R, Pellefigues C, Ronchese F, Lamiable O, Eccles D. Investigation of chimeric reads using the MinION. F1000Research. 2017;

21. Martin S, Leggett RM. Alvis: a tool for contig and read ALignment VISualisation and chimera detection. BMC Bioinformatics. 2021;

22. Marijon P, Chikhi R, Varré JS. Yacrd and fpa: Upstream tools for long-read genome assembly. Bioinformatics. 2020;

23. Xu Y, Lewandowski K, Lumley S, Pullan S, Vipond R, Carroll M, et al. Detection of viral pathogens with multiplex nanopore MinION sequencing: Be careful with cross-Talk. Front Microbiol. 2018;

24. Tvedte ES, Gasser M, Sparklin BC, Michalski J, Hjelmen CE, Johnston JS, et al. Comparison of long-read sequencing technologies in interrogating bacteria and fly genomes. G3 Genes|Genomes|Genetics. 2021;

25. Wick RR, Judd LM, Holt KE. Deepbinner: Demultiplexing barcoded Oxford Nanopore reads with deep convolutional neural networks. PLoS Comput Biol. 2018;

26. Dilthey AT, Jain C, Koren S, Phillippy AM. Strain-level metagenomic assignment and compositional estimation for long reads with MetaMaps. Nat Commun. 2019;

27. Jia B, Xuan L, Cai K, Hu Z, Ma L, Wei C. NeSSM: A Next-Generation Sequencing Simulator for Metagenomics. PLoS One. 2013;

28. Fritz A, Hofmann P, Majda S, Dahms E, Dröge J, Fiedler J, et al. CAMISIM: Simulating metagenomes and microbial communities. Microbiome. 2019;

29. Kolmogorov M, Bickhart DM, Behsaz B, Gurevich A, Rayko M, Shin SB, et al. metaFlye: scalable long-read metagenome assembly using repeat graphs. Nat Methods. 2020;

30. O’Leary NA, Wright MW, Brister JR, Ciufo S, Haddad D, McVeigh R, et al. Reference sequence (RefSeq) database at NCBI: Current status, taxonomic expansion, and functional annotation. Nucleic Acids Res. 2016;

31. Howe KL, Achuthan P, Allen J, Allen J, Alvarez-Jarreta J, Ridwan Amode M, et al. Ensembl 2021. Nucleic Acids Res. 2021;

32. Patro R, Duggal G, Love MI, Irizarry RA, Kingsford C. Salmon provides fast and bias-aware quantification of transcript expression. Nat Methods. 2017;

33. Martin S, Heavens D, Lan Y, Horsfield S, Clark MD, Leggett RM. Nanopore adaptive sampling: a tool for enrichment of low abundance species in metagenomic samples. bioRxiv. 2021;

34. Proctor LM, Creasy HH, Fettweis JM, Lloyd-Price J, Mahurkar A, Zhou W, et al. The Integrative Human Microbiome Project. Nature. 2019;569(7758).

