## Supplementary for "Characterization and simulation of metagenomic nanopore sequencing data with Meta-NanoSim"

**TABLE OF CONTENTS**


**Supplementary methods** 3

**Table S1 Statistical analysis of the abundance estimation on the Adp dataset** 4

**Table S2 MetaFlye assembly comparision** 4

**Fig. S1 Chimera detection and circularity check** 5

**Fig. S2 Average length distribution for each species.** 6

**Fig. S3 Abundance estimation comparison** 7

**Fig. S4 Performance of Meta-NanoSim in simulating one million reads** 8

### Supplementary methods

#### The Expectation-Maximization (EM) algorithm for abundance estimation

**Algorithm 1** EM for metagenome abundance estimation

```
abundance_list = {species1: abundance1; species2: abundance2; …}
base_count = {species1: count1; species2: count2; …}

Start processing uniquely aligned reads:
  for each uniquely aligned read and its source species:
    base_count[species] += aligned bases
  abundance_list = {species: base_count[species]/ sum(base_count[species])}

Start processing multi-aligned reads
while diff >= min(abundance_list.values()) * 0.01:
  E-Step:
    for each multi-aligned read:
      read_abun = the sum of abundances for all possible species for that read
      for each possible species:
        fraction = aligned bases * abundance_list[species] / read_abun
        base_count[species] += fraction
  M-Step:
      abundance_list = {species: base_count[species]/ sum(base_count[species])}
      diff = |abundance_list – prev_abundance_list|
```

**Explanation:**

This EM algorithm estimates the abundance levels in a metagenome from alignments. It first processes uniquely aligned reads to calculate a baseline abundance profile. Then it starts the expectation step, which is to assign multi-aligned bases proportionally to their respective species based on their relative abundances. Next In the maximization step, these multi-aligned bases are used to update the abundance profile. The algorithm then goes back to the expectation step to update the fractions of multi-aligned bases based on the new abundance profile. The E and M steps are alternating until the difference in abundances between two rounds is lower than 1%. Please note that the abundance levels are genomic DNA weight, and it can be used to calculate genome copy number when divided by genome size.

**Table S1 Statistical analysis of the abundance estimation on the *Adp* dataset.**

| Tool | Algorithm | | | | *Adp* dataset | | | |
|---|---|---|---|---|---|---|---|---|
| | E | C | B | R | $R^2$ | Log $R^{2*}$ | Std* | PE |
| Meta-NanoSim | √ | √ | √ | | 0.8831 | 0.9752 | 0.3439 | 2.1020 |
| | | √ | √ | | 0.8831 | 0.9752 | 0.3439 | 2.1020 |
| | √ | | √ | | 0.8830 | 0.9752 | 0.3439 | 2.1037 |
| | | | √ | | 0.8830 | 0.9752 | 0.3439 | 2.1037 |
| Salmon | √ | | | √ | 0.1873 | 0.5045 | 1.5379 | 9.3848 |
| MetaMaps | √ | | | √ | 0.8281 | 0.9643 | 0.4130 | 2.0137 |

**$R^2$:** R-squared, **Std**: standard deviation, **PE**: summation of percent error

**E**: EM algorithm, **B**: base-level quantification, **R**: read-level quantification, **C**: chimeric reads detection

* The expected and estimated abundances are log-transformed before calculating R-squared value and standard deviation.

**Table S2 Runtime and Maximum Resident Set Size comparison for the metaFlye assemblies.**

| Dataset | 1 Million | 2 Million | 4 Million | 10 Million |
|---|---|---|---|---|
| **Runtime (hh:mm:ss)** | 1:14:35 | 2:15:33 | 3:34:31 | 7:00:18 |
| **Maximum Resident Set Size (GB)** | 24.61 | 40.39 | 76.38 | 212.19 |

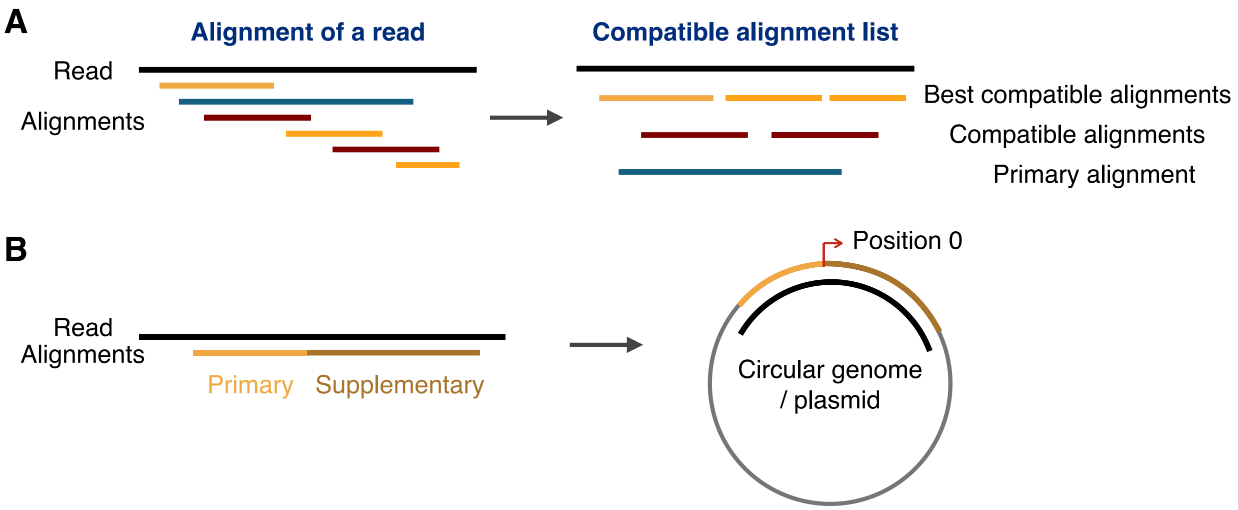


**Fig. S1 Chimera detection and circularity check. A.** Chimeric read detection workflow. Given a read and all its alignments, compatible sub-alignments are computed and sorted to generate a list of compatible alignments. Then the best compatible alignment set is selected based on alignment score and alignment length. If the best compatible alignment set for a read contains two or more compatible alignments, it is considered as chimeric read. **B.** Circularity detection workflow. For circular genomes, if a read surpasses the start position, the alignment is split into a primary and a supplementary alignment. Meta-NanoSim finds these two alignments and treats them together as one alignment.

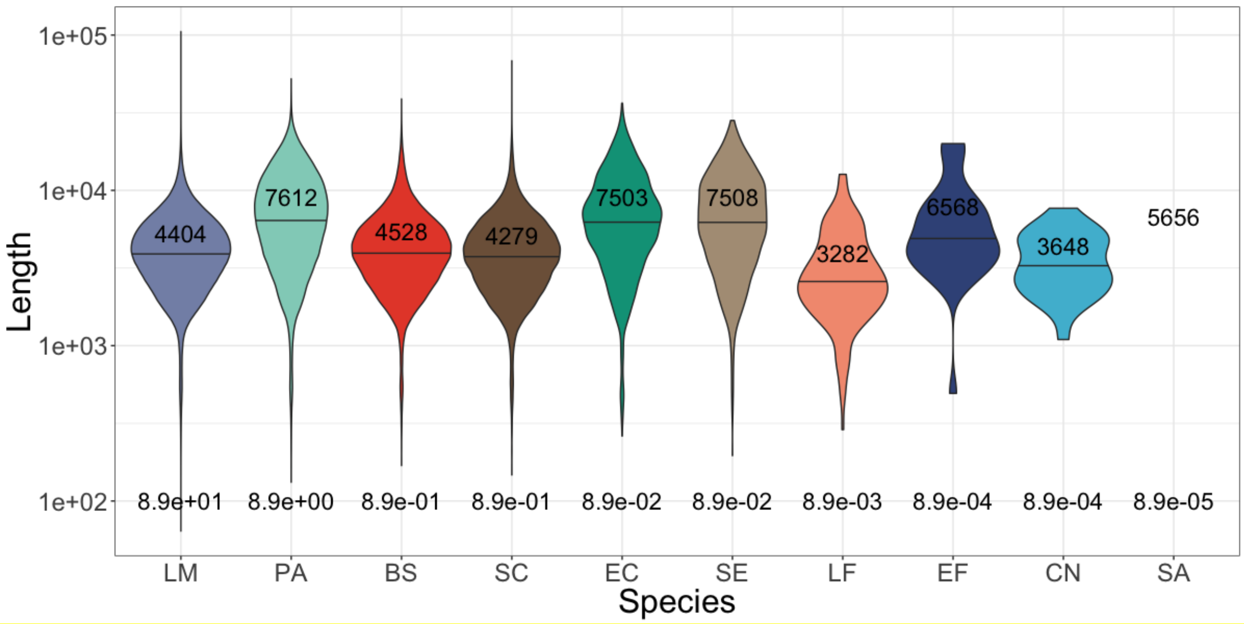


**Fig. S2 Average length distribution for each species.** This plot shows the average read lengths of uniquely aligned reads for each species. The horizontal line in each violin shows the median length and the number above it shows the mean length. The number at the bottom of each violin is the abundance level of each species. There are only two uniquely aligned reads for SA, so there is no violin to show. BS: *Bacillus subtilis*, CN: *Cryptococcus neoformans*, EC: *Escherichia coli*, EF: *Enterococcus faecalis*, LF: *Lactobacillus fermentum*, LM: *Listeria monocytogenes*, PA: *Pseudomonas aeruginosa*, SA: *Staphylococcus aureus*, SC: *Saccharomyces cerevisiae*, SE: *Salmonella enterica*.

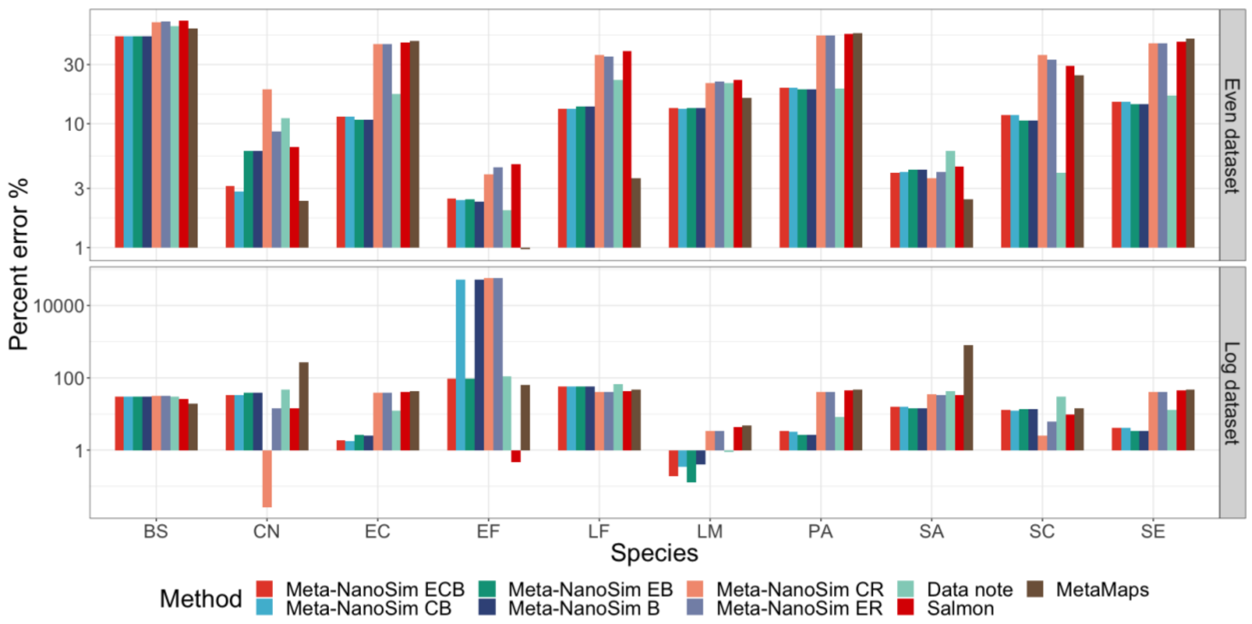


**Fig. S3 Abundance estimation comparison.** The metagenome abundance for Even and Log dataset is calculated using the nine methods, and percent error is calculated as a percentage deviation from the expected values. Meta-NanoSim ECB: EM algorithm + base-level quantification + chimeric reads detection. Meta-NanoSim EB: EM algorithm + base-level. Meta-NanoSim CB: chimeric reads detection + base-level quantification. Meta-Nanosim B: base-level quantification. Meta-NanoSim CR: chimeric reads detection + read-level quantification. Meta-NanoSim ER: EM algorithm + read-level quantification. Salmon: Salmon metagenomic quantification based on Minimap2 alignments. Data note: the abundance reported in the data releasing paper calculated based on Minimap2 alignments. BS: *Bacillus subtilis*, CN: *Cryptococcus neoformans*, EC: *Escherichia coli*, EF: *Enterococcus faecalis*, LF: *Lactobacillus fermentum*, LM: *Listeria monocytogenes*, PA: *Pseudomonas aeruginosa*, SA: *Staphylococcus aureus*, SC: *Saccharomyces cerevisiae*, SE: *Salmonella enterica*.

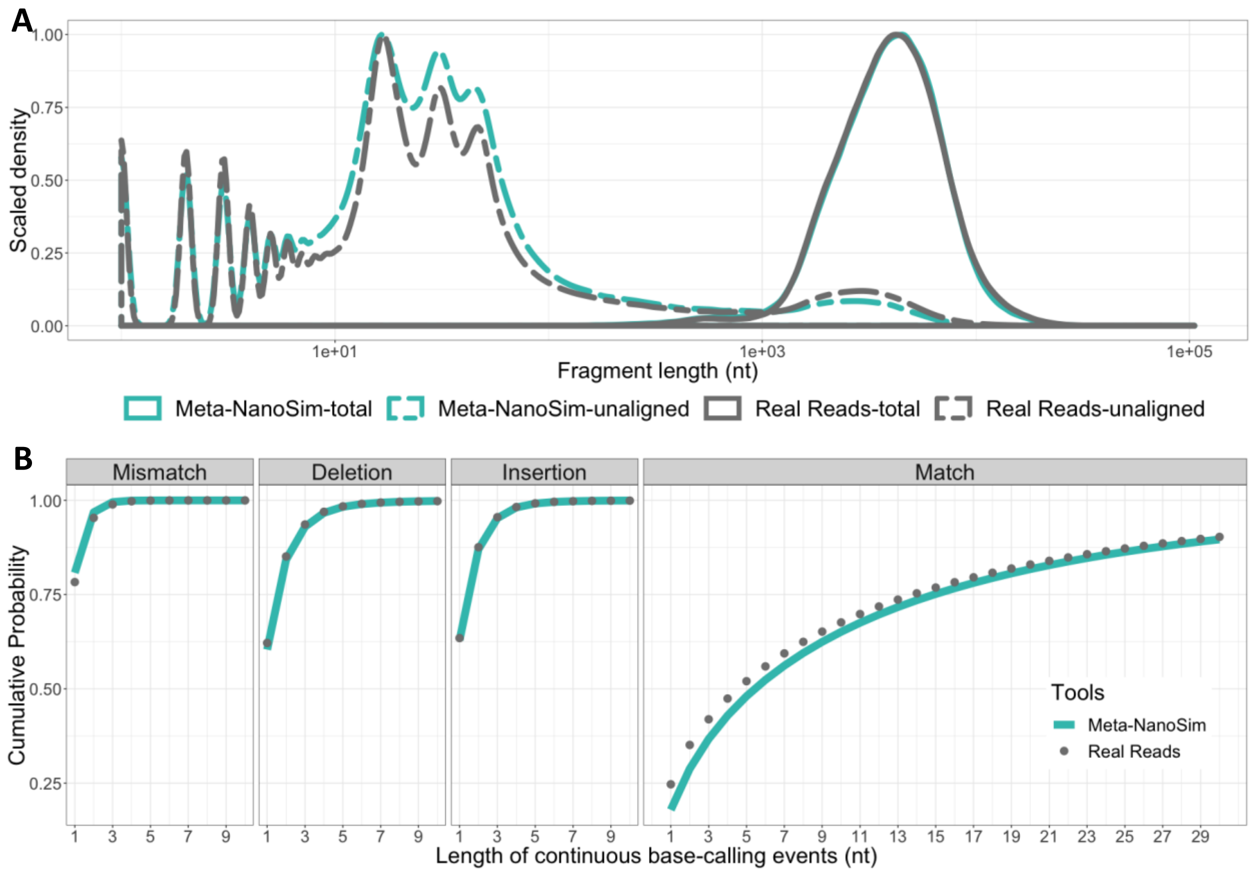


**Fig. S4 Performance of Meta-NanoSim in simulating one million reads.** Meta-NanoSim used the *Log* dataset for simulating the seven-species metagenome (*Adp* dataset). Two samples were simulated at the same time, one million reads in each, with different abundance levels. Details about the simulation can be found in Methods section. **A.** Comparison of read length distributions in the empirical vs. simulated reads. Unaligned length represents the length of unaligned part of each aligned read. **B.** Cumulative probability function of the lengths of matches/errors in empirical and simulated reads.
